# Population-level comparisons of gene regulatory networks modeled on high-throughput single-cell transcriptomics data

**DOI:** 10.1101/2023.01.20.524974

**Authors:** Daniel Osorio, Anna Capasso, S. Gail Eckhardt, Uma Giri, Alexander Somma, Todd M. Pitts, Christopher H. Lieu, Wells A. Messersmith, Stacey M. Bagby, Harinder Singh, Jishnu Das, Nidhi Sahni, S. Stephen Yi, Marieke L. Kuijjer

**Author notes:** Research Computing Group, Boston Children’s Hospital, Harvard Medical School, Boston, MA 02115, USA.

## Abstract

Single-cell technologies enable high-resolution studies of phenotype-defining molecular mechanisms. However, data sparsity and cellular heterogeneity make modeling biological variability across single-cell samples difficult. We present *SCORPION*, a tool that uses a message-passing algorithm to reconstruct comparable gene regulatory networks from single cell/nuclei RNA-seq data that are suitable for population-level comparisons by leveraging the same baseline priors. Using synthetic data, we found that *SCORPION* outperforms 12 other gene regulatory network reconstruction techniques. Using supervised experiments, we show that *SCORPION* can accurately identify differences in regulatory networks between wild-type and transcription factor-perturbed cells. We demonstrate *SCORPION*’s scalability to population-level analyses using a single-cell RNA-seq atlas containing 200,436 cells from colorectal cancer and adjacent healthy tissues. The differences detected by *SCORPION* between tumor regions are consistent across population cohorts, as well as with our understanding of disease progression and elucidate phenotypic regulators that may impact patient survival.

## Introduction

In eukaryotes, gene expression is tightly controlled by the activity of transcription factors, proteins that define cell identity and cellular states by activating or repressing the expression of their target genes in an abundance-dependent manner (1, 2). It is well known that changes in regulatory interactions result in abnormal expression profiles and diseased phenotypes (3–6). Typically, gene regulatory networks are constructed and compared to identify mechanistic alterations in the relationship between transcription factors and their target genes that result in these abnormal phenotypes (7). Transcriptomic data can be used to infer gene regulatory networks by examining the co-expression patterns of genes that are part of the same regulatory programs (8). Depending on the level of detail of the transcriptomic data used to reconstruct the networks, gene regulatory networks can represent the regulatory programs of a specific cell type or the average mechanisms defining the tissue from where the sample was collected (9).

Using the gene expression variability found in RNA-seq data from single cells/nuclei, it is possible to infer the gene regulatory networks for each cell type or cell state within a single sample (10). However, when multiple samples are available, transcriptomes from different samples are typically collapsed by an experimental group before the groups-level comparison is carried out, much like what is done in differential expression analysis (11). In differential network analysis, an aggregate network is built using all of the transcriptomes from each experimental group to represent each one of them (12). Then, in order to learn more about the transcription factor-target gene interactions that support the phenotype of interest, this network is scrutinized or compared to others (13, 14). Although useful, aggregate network models are not designed to account for evaluating between samples’ transcriptional heterogeneity (15).

Pseudo-bulk profiles are frequently calculated in differential gene expression analysis to take into account biological variation between samples (16, 17). However, the biological variability between transcription factors and their target gene interactions must be accurately modeled across multiple samples to identify consistent mechanistic patterns causing phenotypic changes across samples within a population (15, 18, 19). This entails developing time-efficient techniques for constructing highly accurate and comparable gene regulatory networks from single-cell/nuclei RNA-seq data.

Using high-throughput RNA-seq data from single cells or nuclei to create comparable gene regulatory networks is a difficult task (9). This type of data is highly sparse and frequently contains information based on multiple cellular states in a single experiment, making sample comparison difficult (20). Furthermore, non-biological factors frequently affect data during library preparation, reducing our ability to detect biologically accurate correlation structures (21). To address those challenges in differential single-cell gene regulatory network analyses, we present *SCORPION* (<u>S</u>ingle-<u>C</u>ell <u>O</u>riented <u>R</u>econstruction of <u>P</u>ANDA <u>I</u>ndividually <u>O</u>ptimized Gene Regulatory <u>N</u>etworks), a tool that uses coarse-graining of single-cell/nuclei RNA-seq data to reduce sparsity and improve the ability to detect the gene regulatory network’s underlying correlation structure (22). The coarse-grained data generated is then used to reconstruct the gene regulatory network using a network refinement strategy through the PANDA (<u>P</u>assing <u>A</u>ttributes between <u>N</u>etworks for <u>D</u>ata <u>A</u>ssimilation) message passing algorithm (23). This algorithm is designed to integrate multiple sources of information such as protein-protein interaction, gene expression, and sequence motif data to predict accurate regulatory relationships. Thanks to the use of the same baseline priors in each instance, this approach can reconstruct comparable, fully-connected, weighted, and directed transcriptome-wide gene regulatory networks suitable for use in population-level studies.

In this paper, we tested the performance of *SCORPION*’s coarse-grained input data for network modeling using synthetic data via BEELINE, a tool for systematically evaluating cutting-edge algorithms for inferring gene regulatory networks from single-cell transcriptional data (24). We found that networks modeled on data desparsified with *SCORPION* outperform 12 other gene regulatory network reconstruction techniques across seven metrics. Additionally, using supervised experiments, we show that *SCORPION* can precisely identify biological differences in regulatory networks between wild-type cells and cells carrying transcription factor perturbations. Furthermore, we demonstrate *SCORPION*’s scalability to population-level analyses by applying it to a single-cell RNA-seq atlas constructed using publicly available data that includes 200, 436 cells derived from 47 patients and accounts for three different regions of colorectal tumors and healthy adjacent tissue. The differences detected by *SCORPION* between intra- and inter-tumoral regions are consistent with our understanding of disease progression through the chromosomal instability pathway (CIN) that underlies the majority of all colon cancers (25). Findings were confirmed in an independent cohort of patient-derived xenografts from left and right-sided tumors and provide insight into the regulators associated with the phenotypes and the differences in their survival rate.

## Results

### The *SCORPION* algorithm

*SCORPION* is an R package that generates through five iterative steps comparable, fully connected, weighted and directed transcriptome-wide gene regulatory networks from single-cell transcriptomic data that are suitable for their use in population-level studies (Figure 1A). To begin, the highly-sparse high-throughput single-cell/nuclei RNA-seq data is coarse-grained by collapsing a *k* number of more similar cells identified at the low dimensional representation of the multidimensional RNA-seq data (26). This approach reduces sample size while also decreasing data sparsity, allowing us better to capture the strength of the relationship between genes’ expression (22).

The second step is to construct three distinct initial unrefined networks, as described in the PANDA algorithm: the co-regulatory network, the cooperative network, and the regulatory network (23). The co-regulatory network represents the co-expression patterns between genes. This network is constructed using correlation analyses over the generated coarse-grained transcriptomic data. The cooperative network accounts for the known protein-protein interactions between transcription factors. This information is downloaded from the STRING database (27). The third network is the unrefined regulatory network that describes the relationship between transcription factors and their target genes through transcription factors footprint motifs found in the promoter region of each gene (28).

Following the construction of the three networks, a modified version of the Tanimoto similarity designed to account for continuous values is used to generate the availability network (*A*_*ij*_), representing the information flow from a transcription factor *i* to a gene *j*, describing the accumulated evidence for how strongly the transcription factor influences the expression level of that gene, taking into account the behavior of other genes potentially targeted by that transcription factor. In addition, the responsibility network (*R*_*ij*_) is generated by computing the similarity between the cooperativity network and the regulatory network. The responsibility represents the information flowing from a transcription factor *i* to a gene *j* and captures the accumulated evidence for how strongly the gene *j* is influenced by the activity of that specific transcription factor, taking into account other potential regulators of gene *j*.

The average of the accessibility and the responsibility networks is computed in the fourth step, and the regulatory network is updated to include a user defined proportion (*α* = 0.1 by default) of the information provided by the other two original unrefined networks. The cooperativity and co-regulatory networks are also updated in the fifth step using the new information contained in the updated regulatory network. Steps three through five are repeated iteratively until the hamming distance between the networks reaches a user-defined threshold (0.001 by default). When convergence is reached, the refined regulatory network is returned as a matrix with transcription factors in the rows and target genes in the columns. The matrix values encode the strength of the relationship between each transcription factor and gene.

### *SCORPION* outperforms 12 other algorithms for single– cell gene regulatory network construction

To provide a comparison of how data desparsification in *SCORPION* would affect downstream network modeling, we tested its performance to that of other algorithms. To do so, we conducted a systematic comparison of network construction algorithms using BEELINE, an evaluation tool designed for this purpose (24). *SCORPION* was tested and compared to 12 different algorithms (9, 29–39). Each method’s performance in recovering gene-to-gene relationships was compared to ground-truth interactions between genes generated using pre-set parameters without other information than the expression matrix. According to our findings, *SCORPION* generates 18.75% more precise (higher precision) and sensitive (higher recall) single-cell gene regulatory networks than other methods. We also found that, on average, *SCORPION* ranks first when compared to other methods using seven different metrics related to network construction (Figure 1B, Supplementary Table S1).

**Fig. 1.**
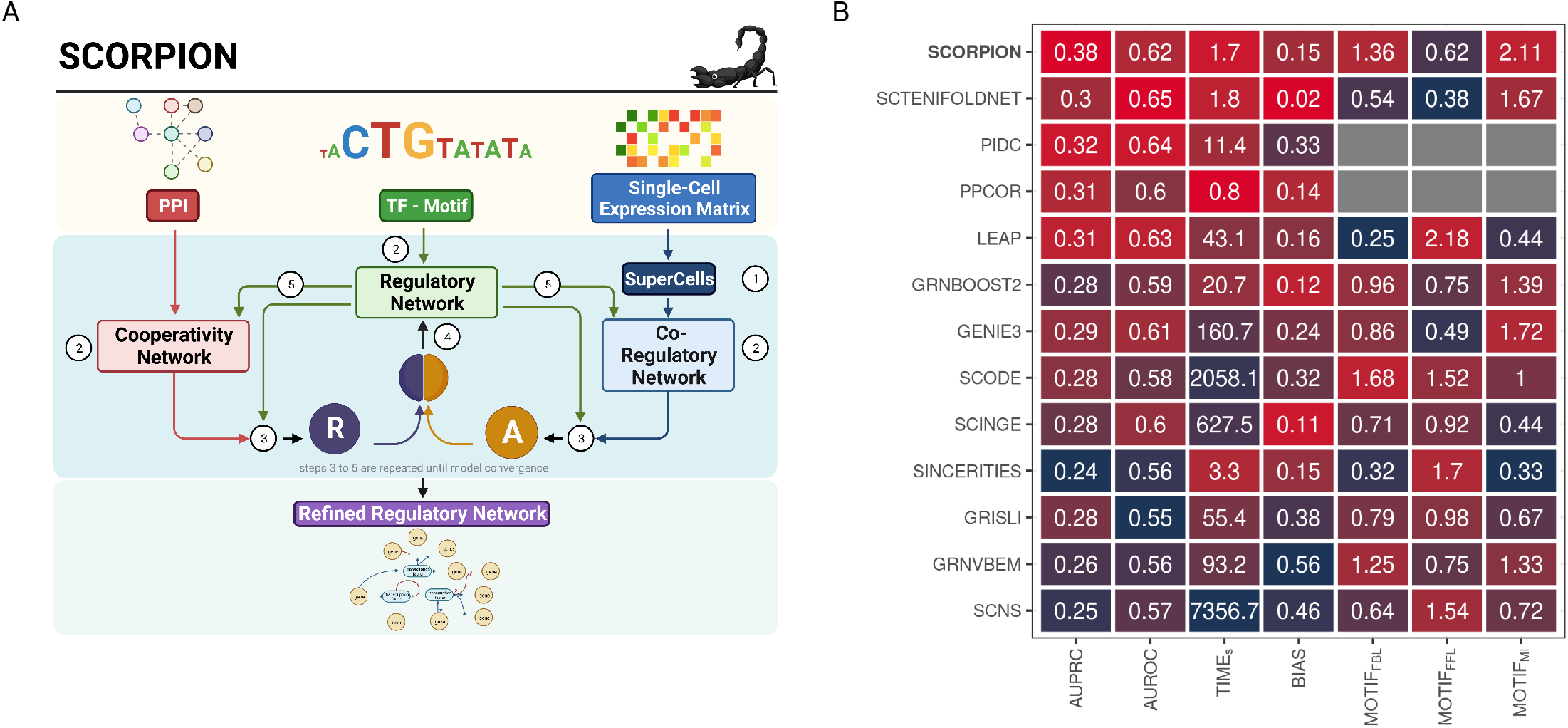
Overview and benchmarking of desparsification with *SCORPION*. **(A)** *SCORPION* uses the PANDA message passing algorithm to integrate data from multiple sources, including protein-protein interactions, single-cell gene expression, and sequence motif data, to predict accurate regulatory relationships. In five iterative steps, *SCORPION* generates comparable, fully-connected, weighted, and directed transcriptome-wide gene regulatory networks from from single-cell transcriptomic data suitable for use in population-level studies. **(B)** The performance of 13 single-cell gene regulatory network construction methods was evaluated using BEELINE and the same curated synthetic dataset. Methods are ranked based on their average performance across seven different metrics. If the metric was not quantifiable, gray squares are displayed. Performance in each metric is color coded from red (best) to blue (worst). The acronyms of the metrics are explained in Methods.

The curated dataset provided by BEELINE to perform the benchmark of the different tools is much simpler than the transcriptome-wide gene regulatory network required in reality to identify mechanistic changes in gene regulation that support the observed phenotypes. In fact, it is known that incorporating prior information on TF binding into regulatory network reconstruction algorithms improves predictions of regulation (40). For that reason, after having tested the outperformance of *SCORPION*’s desparsification approach on synthetic data, we chose to apply the complete *SCORPION* framework (desparsification with Super-Cells and message passing between prior regulatory, cooperativity, and co-regulatory networks) directly to curated real datasets and assess the biological relevance of the generated gene regulatory networks.

### *SCORPION* accurately detects changes in transcription factor activity and their impact on target genes

We used two curated real datasets generated using 10× Genomics’ high-throughput single-cell/nuclei RNA-seq technologies to evaluate *SCORPION*’s performance in identifying changes in transcription factor activity and their impact on target genes. The first dataset was generated to examine the redundant effect of *Hnf4α* and *Hnf4γ* transcription factors in the intestinal epithelium of mice through a double knockout (DKO) experiment (41). The second dataset was designed to investigate the role of over-expressing the *DUX4* transcription factor on human embrionic stem cells (hESCs) during human zygotic genome activation-like transcription *in-vitro* (42).

For the first dataset, two independent single-cell gene regulatory networks were built to model the regulatory mechanisms on *Hnf4αγ*^WT^ (wild-type, *n* = 4, 100) and *Hnf4αγ*^DKO^ (*n* = 4, 200) mice intestinal epithelial cells. The *Hnf4αγ*^WT^ network models the regulation of 4, 255 genes by 603 transcription factors, while the *Hnf4αγ*^DKO^ network account for the regulation of 3, 384 genes by the same amount of transcription factors as in *Hnf4αγ*^WT^. We used the subnetwork representing the regulatory mechanisms of the 2, 990 genes that overlapped in both networks for comparison. We focused on the differences in the outdegrees (the sum of edge weights from a TF to all genes) of the *Hnf4α* and *Hnf4γ* transcription factors because they represent the changes on the transcription factor’s activity over their target genes’ expression after perturbation. In both cases, we observed a shift in the weights of the links between the perturbed transcription factors and their target genes (Figures 2A and 2E). The paired weight differences were found to be highly significant (*t*-test, *P* = 1.1 × 10^*−*85^), and the direction of the shift (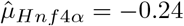 and 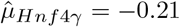) consistent with the perturbation targeted (downregulation) in the cells during the experimental design (Figures 2B and 2F).

**Fig. 2.**
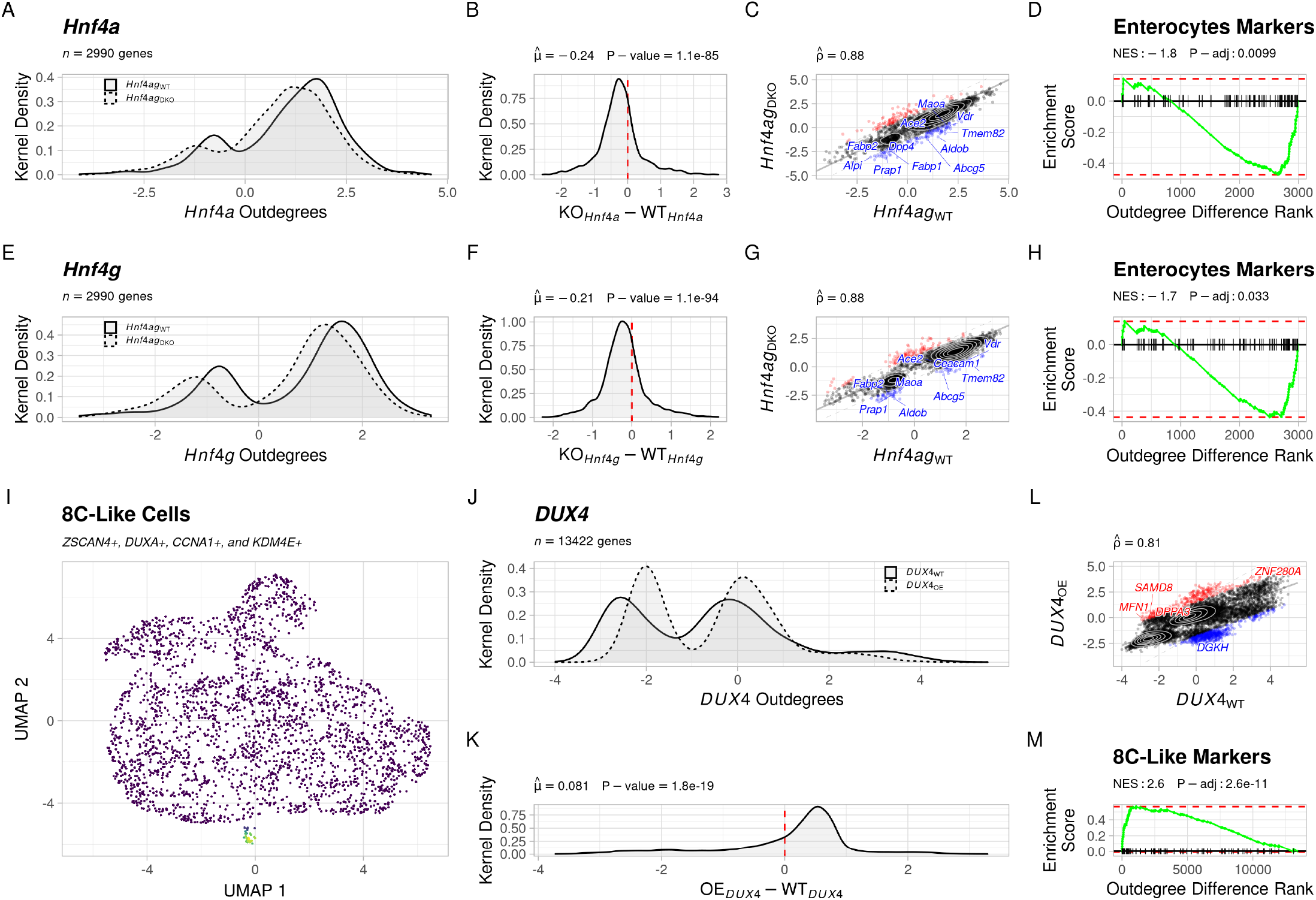
Evaluation of *SCORPION* ‘s ability to detect changes in transcription factor activity and their impact on target genes. **(A)** Differences in the distribution of the outdegree weights for the *Hnf4α* transcription factor in *Hnf4αγ*^WT^ and *Hnf4αγ*^DKO^ mice intestinal epithelium cells. **(B)** Distribution of the paired differences between the outdegrees of the *Hnf4α* transcription factor (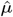 and P-value were calculated using a one-sample *t* -test). **(C)** Spearman correlation 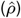 of the outdegrees for the *Hnf4α* transcription factor in *Hnf4αγ*^WT^ and *Hnf4αγ*^DKO^ mice intestinal epithelium cells. Genes out of the 95% confidence interval are color coded and labeled (in red if up-regulated, and in blue if down-regulated). **(D)** Gene set enrichment analysis of Enterocytes marker genes using the paired differences between the outdegrees of the *Hnf4α* transcription factor. **(E)** Differences in the distribution of the outdegrees for the *Hnf4γ* transcription factor in *Hnf4αγ*^WT^ and Hnf4*αγ*^DKO^ mice intestinal epithelium cells. **(F)** Distribution of the paired differences between the outdegrees of the *Hnf4γ* transcription factor (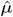 and P-value were calculated using a one-sample *t* -test). **(G)** Spearman correlation 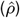 of the outdegrees for the *Hnf4γ* transcription factor in *Hnf4αγ*^WT^ and *Hnf4αγ*^DKO^ mice intestinal epithelium cells. Genes out of the 95% confidence interval are color coded and labeled (in red if up-regulated, and in blue if down-regulated). **(H)** Gene set enrichment analysis of the Enterocytes marker genes using the paired differences between the outdegrees of the *Hnf4γ* transcription factor. **(I)** Uniform Manifold Approximation and Projection (UMAP) of hESCs. 8C-cell-like cells are highligthed. **(J)** Differences in the distribution of the outdegrees for the *DUX4* transcription factor in *DUX4*^WT^ and *DUX4*^OE^ hESCs. **(K)** Distribution of the paired differences between the outdegrees of the *DUX4* transcription factor (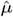 and P-value were calculated using a one-sample *t* -test). **(L)** Spearman correlation 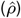 of the outdegrees for the *DUX4* transcription factor in *DUX4*^WT^ and *DUX4*^OE^ hESCs. Genes out of the 95% confidence interval are color coded and labeled (in red if up-regulated, and in blue if down-regulated). **(M)** Gene set enrichment analysis of the 8C-like cells marker genes using the paired differences between the outdegrees of the *DUX4* transcription factor.

We found 221 and 211 large changes (out of the 95% confidence interval, 181 genes shared, Jaccard Index = 0.819) after perturbation in *Hnf4α* and *Hnf4γ* outdegrees, respectively. These changes (Figures 2C and 2G) highlight 84 shared genes with decreased activation signal (downregulation) from 114 in *Hnf4α* and 95 in *Hnf4γ* (Jaccard Index = 0.672), as well as 97 shared genes with increased activation signal (upregulation) from 107 in *Hnf4α* and 116 in *Hnf4γ* (Jaccard Index = 0.769). The high overlap (81.9%) in the top-most perturbed target genes discovered after the double knockout supports the paralog redundant activity of *Hnf4α* and *Hnf4γ* in the intestinal epithelium of mice. Additionally, in agreement with what the dataset’s original authors reported (41), when we performed gene set enrichment analysis using the paired differences between the weights of the link between the transcription factors and their target genes, we found that *Hnf4α* and *Hnf4γ* perturbations have a significant (NES *<* 0, and *FDR <* 0.05) impact on reducing the expression of the canonical marker genes associated with enterocyte’s identity development (Figures 2D and 2H, Supplementary Table S2 and S3).

For the second dataset, as before, we constructed two independent gene regulatory networks to model the regulatory mechanisms on wild-type human embryonic stem cells (hESCs) and the effect of over-expressing (OE) the *DUX4* transcription factor on them. The resulting two gene regulatory networks represent the regulatory effect of 622 transcription factors over 13, 422 genes in 970 *DUX4*^*W T*^ hESCs and a subset of 55 *DUX4*^*OE*^ hESCs exhibiting the canonical marker genes (*ZSCAN4, DUXA, CCNA1*, and *KDM4E*) of 8C-like cells (Figure 2I, Supplementary Table S4). When we compared the transcription factor activity of *DUX4* in both networks, we noticed a shift in distribution of the weights of the links before and after the transcription factor was overexpressed (Figure 2J). In agreement with the experimental design targeted in the hESCs, we found that the paired differences in the weights of the links between *DUX4* and its target genes are significantly (*t*-test, P *<* 0.0001) shifted to the positive side (Figure 2K), inducing upregulation of its target genes. We found 999 extreme link weight changes out of the 95% confidence interval, that represent 624 and 375 target genes down- and up-regulations associated with the overexpression of *DUX4* on hESCs respectively (Figure 2L). When we performed gene set enrichment analysis using the paired differences between the weights of the links between *DUX4* and its target genes, we found that these are positively associated (NES *>* 0, *P <* 0.05) with the overexpression of highly expressed genes in 8C-like cells such as *ADD3, ALPG, BCAT1, DPPA3, EXOSC10, HIPK3, NEAT1, ODC1, RBBP6, RBM25, SAMD8, SLC2A3, WDR47*, and *ZNF217* (Figure 2M, Supplementary Table S5).

These findings confirm that *SCORPION* can detect experimentally targeted changes in transcription factor activity and represent the impact of those changes on the resulting gene regulatory networks. This holds true when comparing two networks. However, since *SCORPION* networks are refined using a message-passing algorithm, the only difference between the resulting networks is given by the correlation structure provided by the RNA-seq data from single cells/nuclei used to generate the co-regulatory network. This feature, in conjunction with the short time of construction (Figure 1B), makes *SCORPION* suitable for the generation of comparable gene regulatory networks in a pipeline scalable to population-level studies targeting the identification of differences in gene regulation. In order to showcase this feature, we chose to use *SCORPION* to reconstruct gene regulatory networks for each cell type within each sample in a multi-sample single-cell atlas of colorectal cancer that includes cells from both nearby normal tissue and three distinct tumor regions.

### *SCORPION* gene regulatory networks recapitulate cellular identity and disease status

We generated a multisample single-cell RNA-seq atlas containing the transcriptomes of cells from adjacent healthy tissue and three different regions of colorectal tumors, including metastasic, core, and border tissue aiming to characterize the regulatory mechanisms driving the development and progression of colorectal cancer. To begin, we gathered single-cell RNA-seq data from five publicly available datasets comprising 303, 221 cells derived from 47 donors. After quality control, 200, 439 were kept on the study (Figure 3, Panels A-D; Supplementary Table S6). *SCORPION* was then used to generate a gene regulatory network for each cell type (with at least 30 cells) within each sample included in the atlas after cells were annotated using canonical markers (Supplementary Figure S1). At the end, we generated 560 transcriptome-wide gene regulatory networks that account for the regulatory effect of 622 transcription factors over 17, 425 target genes (a total of 10, 838, 350 links) in each network.

**Fig. 3.**
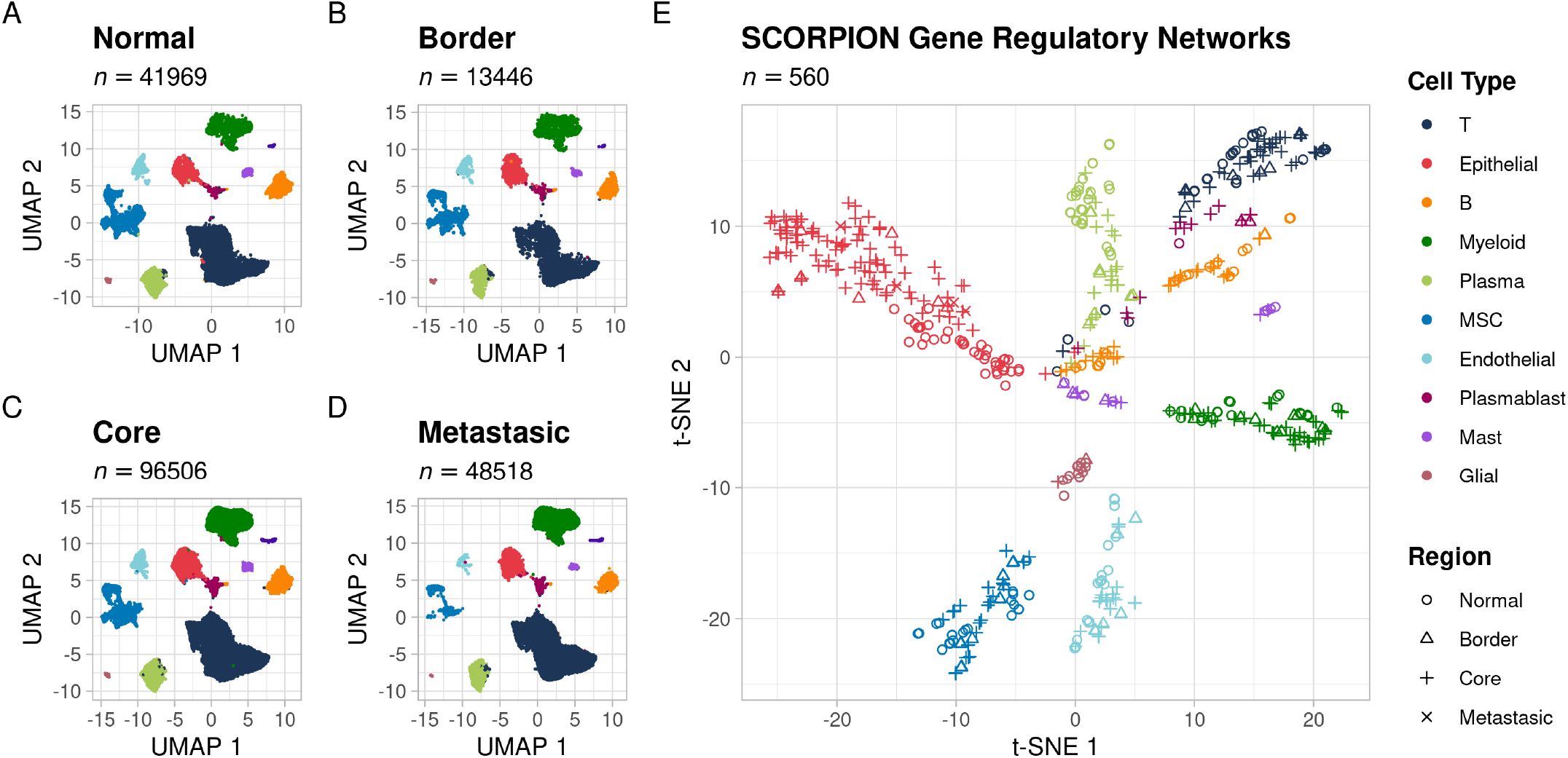
Low-dimensional representation of transcriptomes and gene regulatory networks from colorectal cancer and adjacent healthy tissue. **(A)** UMAP of cells from healthy adjacent tissue. **(B)** UMAP of cells from tumor border tissue. **(C)** UMAP of cells from tumor core tissue. **(D)** UMAP of cells from liver metastasic tissue. **(E)** t-SNE of gene regulatory networks from colorectal cancer and adjacent healthy tissue generated by *SCORPION*.

We used the network’s indegrees (the sum of the weights from all transcription factors to a gene) to generate a t-distributed stochastic neighbor embedding (t-SNE) low-dimensional representation of the information contained in the networks. We found that networks of cells of the same type cluster together regardless of tissue of origin (Figure 3E). This reaffirms *SCORPION*’s ability to accurately identify the differences in regulatory mechanisms defining cell type identity across multiple samples.

We chose cells from the core tissue, border tissue, and adjacent healthy tissue from four different donors to compare their similarities in order to assess the reproducibility of the built gene regulatory networks (Supplementary Figure S2). We found that, on average, the similarity between the cancer tissue (core and border) is significantly (*t*-test, *P* = 3.5 × 10^*−*3^) higher (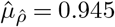, Supplementary Figure S2A) than the one observed when compared the cancer tissue with the healthy adjacent one (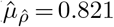, Supplementary Figure S2B). This outcome confirms our previous findings, in which we were able to reconstruct two gene regulatory networks that represented the control of 15, 493 genes through 622 transcription factors in T-cells derived from two samples taken from the same benign polyp in a female donor with adenomatous polyposis. Those networks exhibited a highly positive and significant Spearman correlation coefficient (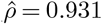, *P* = 2.2 *×* 10^*−*16^) (43, 44).

### *SCORPION* gene regulatory networks reveal patterns of colorectal cancer progression and sidedness

One of the most significant advantages of using single-cell/nuclei RNA-seq data is the ability to characterize the molecular mechanisms underlying disease at the cell-type specific level (2). Because colorectal cancer is an epithelial cancer, we decided to focus on the molecular mechanisms that drive disease progression in epithelial cells. We selected the 149 single-cell gene regulatory networks generated for this cell type among the four tissues (healthy *n* = 42, border *n* = 9, core *n* = 94, and metastasis *n* = 4), and used linear regression to investigate each of the 9, 532, 150 links between 622 transcription factors and 15, 325 target genes aiming to identify linear patterns of up- or down-regulation across these links. Our reasoning was that healthy adjacent tissue (encoded as 1) is transitionally transformed into malignant tissue along the border (encoded as 2), and disease signals will be increased in the tumor’s core (encoded as 3), and metastasic tissue (encoded as 4). We calculated a *β* coefficient and associated adjusted for multiple testing P-value for each link (Figure 4A). We found 5, 202, 588 links with a absolute value of *β* greater than 0 and a false discovery rate less than 0.05 (45). We treated these *β* coefficients as weights in the generated network representing colorectal cancer progression (Figure 5, Supplementary Table S7).

**Fig. 4.**
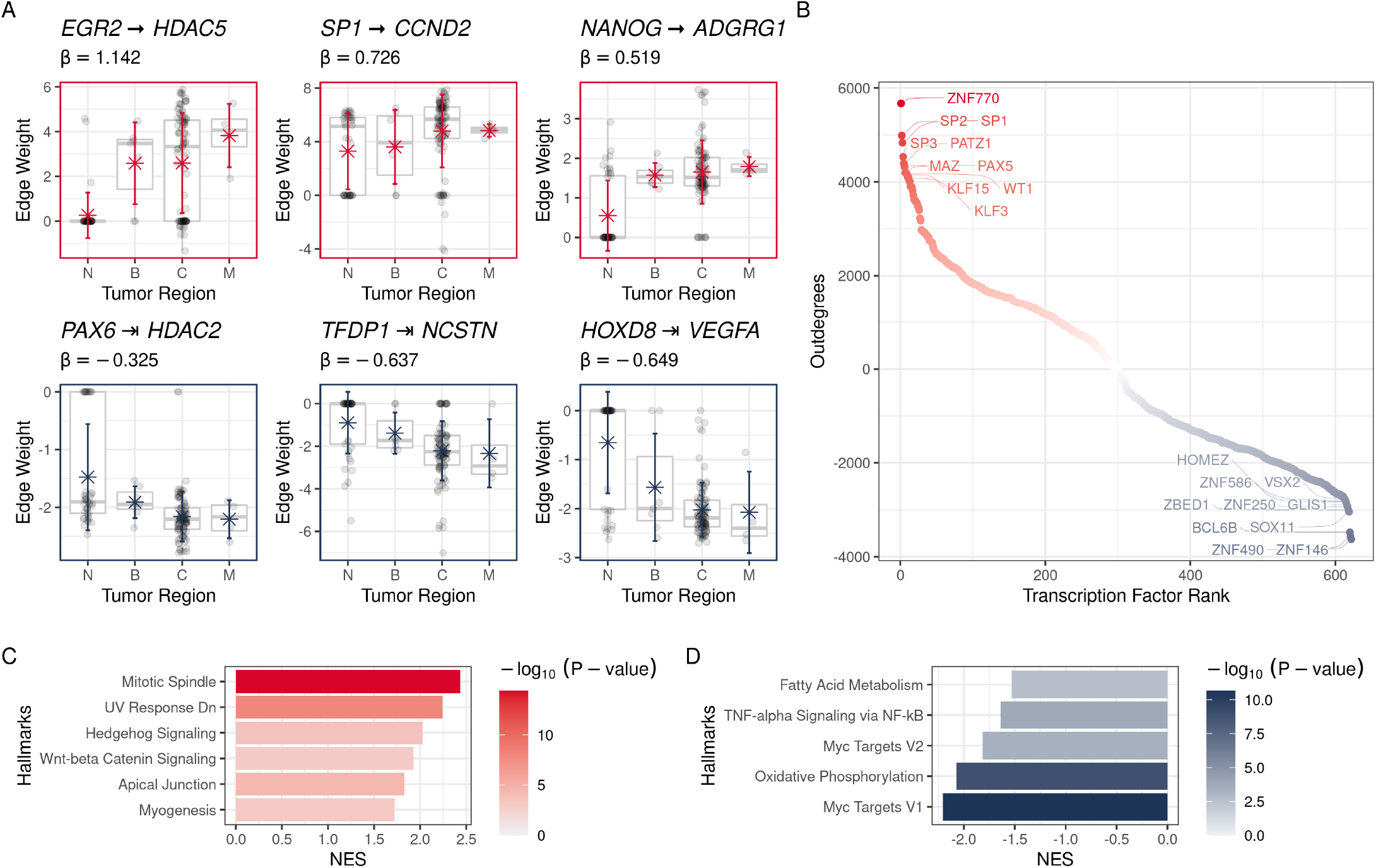
Differential network analysis of epithelial cells during colorectal cancer progression. N: Adjacent normal tissues, B: Border of the tumor, C: Core of the tumor, M: Liver Metastases. **(A)** Examples of significant interactions between transcription factors and target genes linearly increasing or decreasing during colorectal cancer progression. **(B)** Ranked list of transcription factors based on the transcription factor activity in the gene regulatory network illustrating the progression of colorectal cancer. **(C)** Significantly upregulated hallmarks found in the gene regulatory network illustrating the progression of colorectal cancer. **(D)** Downregulated hallmarks found in the gene regulatory network illustrating the progression of colorectal cancer.

**Fig. 5.**
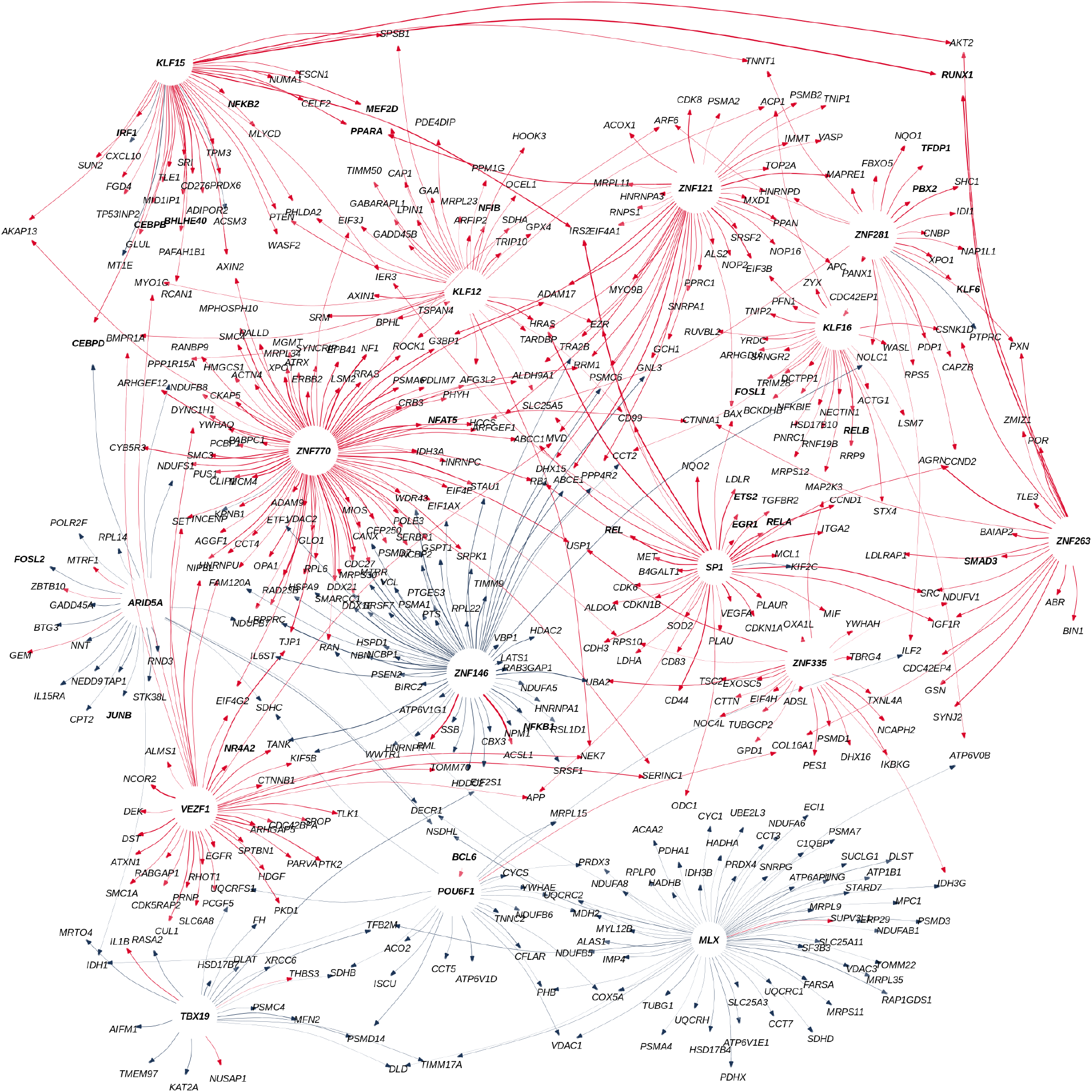
Gene regulatory network illustrating the progression of colorectal cancer. Transcription factors with the highest activities up- or down-regulated are displayed in bold letters. The graph’s edges are color-coded in red for up-regulated and blue for down-regulated interactions. The width of the displayed edge encodes the magnitude of the computed beta coefficient.

We found that some of the identified interactions have directions that are consistent with previously reported oncogenic transformation patterns necessary for the growth and development of colorectal tumors (Figure 4A). For example, upregulation of *EGR2* is required for colon cancer stem cells survival and tumor growth (46), upregulation of *HDAC5* promotes colorectal cancer cell proliferation (47), upregulation of *SP1* activate the Wnt-beta catenin pathway in colorectal cancer (48), upregulation of *CCND2* in conjunction with *JAK2* and *STAT3* promotes colorectal cancer stem cell persistence (49), upregulation of *NANOG* modulates stemness in human colorectal cancer (50), upregulation of *ADGRG1* promotes proliferation of colorectal cancer cells and enhances metastasis via epithelial to mesenchymal transition (51). Examples, where edge weights are reduced through tumor progression, include the inhibition of the epithelial to mesenchymal transition during cancer metastasis by *HDAC2* (52), and the tumor-suppressing role in colorectal cancer by *HOXD8* that act as an apoptotic inducer (53).

To identify the major drivers of colorectal cancer progression, we calculated transcription factor overall association as the (outdegree) sum of all the beta coefficients for each transcription factor and their target genes. We found that the top ten most associated transcription factors across colorectal cancer development are *ZNF770, SP1, SP2, SP3, PATZ1, MAZ, PAX5, KLF15, WT1*, and *KLF3*. Among these, *SP1, WT1, PAX5* and *KLF3* are known to be associated with transcriptional misregulation in cancer (Hyper-geometric test, KEGG database, *OR* = 70.22, *FDR <* 0.0001). On the other hand, the top ten associated transcription factors with reduced outdegrees throughout tumor progression are *ZNF146, ZNF490, BCL6B, SOX11, ZBED1, ZNF250, GLIS1, ZNF586, HOMEZ*, and *VSX2* (Figure 4B).

We also calculated the network’s indegrees by aggregating the regulation of all transcription factors over a target gene. We used this vector of aggregated weights representing the rate of change of each edge during disease progression after performing linear regression of the indegrees to evaluate gene set enrichment using the hallmarks of cancer as reference (54, 55). Out of the 50 hallmarks, we found 11 significantly (*FDR <* 0.05) perturbed. *Mitotic spindle, Hedgehog signaling*, and *Wnt-beta catenin signaling* were among the six hallmarks found to be upregulated (NES *>* 0). These three characteristics are part of a well-known colorectal cancer pathway known as the chromosomal instability (CIN) pathway (25). The CIN pathway is linked to an increase in genomic instability, which is critical for the development of colorectal cancer. CIN is also the most common cause of colorectal cancer (56). Additionally, we found that the c-Myc pathway in the epithelial cells of the tumor’s core and metastasis regions was significantly downregulated (NES *<* 0). This is in line with earlier reports suggesting that low c-*MYC* levels enable cancer cells to survive in the presence of low levels of oxygen and glucose, which are characteristic of the tumor’s core (57).

Overall, we found that the regulatory patterns represented in the gene regulatory networks generated by *SCORPION* to characterize the progression of colorectal cancer in epithelial cells strongly agree with our understanding of the disease’s progression. These high-quality data with unparalleled resolution due to the use of single-cell RNA-seq show that *SCORPION* is suited for the construction of comparable gene regulatory networks to support population-level comparisons aimed at identifying differences in gene regulation.

We next wanted to demonstrate the potential of *SCORPION* to identify differences in gene regulatory networks between conditions. There are four accepted consensus molecular categories for colorectal cancer, CMS1 (microsatellite instability immune), CMS2 (canonical), CMS3 (metabolic), and CMS4 (mesenchymal), which were determined based on the tumor’s composition and mutational status (58, 59). A genetic cascade of changes causes the normal colonic epithelium to first become an adenoma and subsequently an adenocarcinoma as colorectal cancer progresses (60). For this reason, it is essential to first comprehend and give priority to the regulatory mechanisms of malignant epithelial cells in order to develop pharmacological options for patients. It is well recognized that the origin, phenotype, and prognosis of cancer arising from different sides of patients’ intestines vary (61). Whereas differences in tumor composition and differential gene expression at the single-cell atlas level have been reported before (62), a differential gene regulatory network analysis aiming to identify regulatory drivers of the differences has never been conducted at this level of resolution. We therefore chose to contrast the regulatory processes defining colorectal tumors arising on the left (splenic flexure, sigmoid colon, descending colon and rectum) and right (cecum, apendix, ascending colon and hepatic flexure) sides of the patients’ intestines.

To determine the drivers of regulatory differences across epithelial cells from the core of 11 right-sided and 22 left-sided colorectal tumors (see Methods), we computed transcription factor targeting (outdegree) for each of the 622 transcription factors in each network independently (Figure 6A). After comparing the two groups, we found 118 transcription factors with enhanced activity in right-sided colorectal cancer in contrast with the 287 found with enhanced targeting in left-sided colorectal cancer (Figure 6B). Among the top ten more active transcription factors in left-sided colorectal cancer (Figure 6C) we found a significant enrichment of transcription factors associated with unfolded protein response (*NFYA* and *CEBPG*, Hypergeometric test, *FDR <* 0.01). In right-sided ones (Figure 6D), we found an enrichment of transcription factors associated with TNF-alpha signaling via NF-kB (*KLF9, NFKB1* and *NFKB2*, Hypergeometric test, FDR *<* 0.001). A thorough examination of the unfolded protein response and the NF-kB signaling pathways in colorectal cancer has previously been reported (63, 64). We found that the most significant drivers of the differences between left-sided and right-sided colorectal cancer found in our analysis are *ZNF350* (*t*-test, *FDR* = 0.024) and *NFKB2* (*t*-test, *FDR* = 0.032) respectively.

**Fig. 6.**
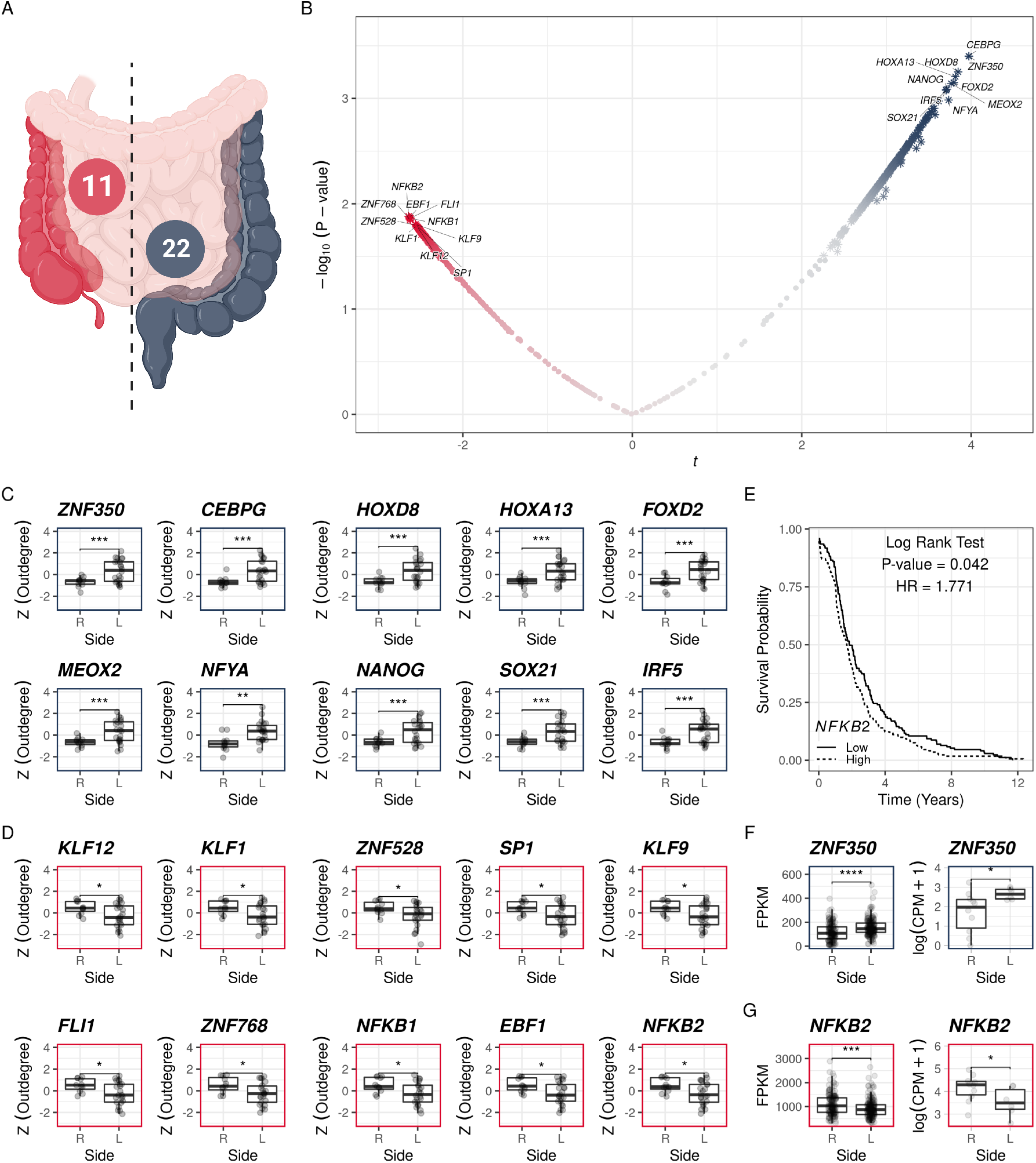
Regulatory differences between right-sided and left-sided colorectal cancer epithelial cells. **(A)** A diagram depicting the left and right side of the intestines. The number of samples for each group is also shown. **(B)** Volcano plot displaying the differences in transcription factor activity between right-sided and left-sided colorectal cancer epithelial cells. **(C)** Top 10 most active transcription factors in epithelial cells from left-sided colorectal cancer. **(D)** Top 10 most active transcription factors in epithelial cells from right-sided colorectal cancer. **(E)** Differences in patient survival rates according to *NFKB2* expression in patients with primary colorectal cancer. **(F)** Consistent differences in gene expression for the *ZND350* transcription factor in the TCGA data and our own dataset. **(G)** Consistent differences in gene expression for the *NFKB2* transcription factor in two independent patient cohorts. *t* –test P-value significance codes: * ≤ 0.05, ** ≤ 0.01, *** ≤ 0.001, ****≤ 0.0001

When these two patterns are combined, they are consistent with the significantly worse survival rate of patients with right-sided colorectal malignancies (65). The methylation of the *ZNF350* transcription factor’s promoter region, which causes its downregulation, is known to stimulate colon cancer cell migration (66). Additionally, over-expression of *NFKB2* is a known prognostic marker of poor survival in colorectal cancer (67). To cross-validate these relationships, we first compared the averaged survival rates based on *NFKB2* expression in patients with primary tumors in the cecum, apendix, ascending colon, hepatic flexure, splenic flexure, sigmoid colon, descending colon, and rectum from the TCGA-COAD and TCGA-READ projects (68). We confirmed the association between the level of *NFKB2* expression and the average survival rate of the patients (Log-rank test, *P* = 0.042, Figure 6E). Following that, we compared the levels of expression of the two transcription factors in primary colorectal tumors on the left and right sides of the intestine. We found that, in both cases, the patterns identified by *SCORPION* and represented in the gene regulatory networks are consistent in directionality and significance with the level of expression observed in the primary tumors from the TCGA data (Left panels in Figures 6F and 6G).

To further cross-validate our findings and assess the reliability of this pattern in a smaller population, we compared the expression levels of both transcription factors in a new dataset of 15 patient-derived xenograft models (PDXs, see Methods) generated by us (Supplementary Table S8). Nine samples were from right-sided and six from left-sided colorectal tumors. Here, as before with the TCGA data, we demonstrated that the patterns identified by

*SCORPION* and represented in the gene regulatory networks are consistent in both directionality and significance with the level of expression observed (Right panels in Figures 6F and 6G). These consistent findings in three independent datasets support the ability of *SCORPION* to identify reliable transcription factor differential activities driving observed phenotypes in a cell-type specific manner and highlight potential targets for developing pharmacological options for patients with right-sided colorectal cancer aiming to improve their poor survival rate.

These findings highlight *SCORPION*’s ability to identify not only intra-tumoral characteristics affecting patient survival but also novel biomarkers and appropriate targets for developing pharmacological options for patients.

## Discussion

We present *SCORPION*, a tool for constructing fully connected, weighted, and directed transcriptome-wide gene regulatory networks from single-cell transcriptomic data that can be used in population-level studies. Despite the fact that there are numerous methods for constructing gene regulatory networks from single-cell transcriptomes, these do not allow for comparative analysis while accounting for the heterogeneity between samples of the regulatory mechanisms defining a phenotype at the population level.

*SCORPION* represents a breakthrough in gene regulatory network construction using single-cell transcriptomics. *SCORPION* integrates multiple sources of regulatory information as a baseline and refines this knowledge using an optimization algorithm to produce gene regulatory networks that account not only for the activity of the transcription factors but also for their cooperativity with high accuracy. The use of data other than gene expression distinguishes *SCORPION* from most other methodologies (9, 29–39). Compared with other algorithms that do incorporate prior information on transcrition factors, such as SCENIC (8) and SCIRA (69), *SCORPION* uses the information about the motif footprints during the construction of the network and not only to characterize the activity of the transcription factors. Furthermore, unlike SCENIC, *SCORPION* employs an association metric (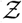 -scores) with a defined underlying distribution (𝒩) that make possible the comparison of weights across experiments and allowed us to identify edges associated with colorectal cancer progression, and, like SCIRA, *SCORPION* allows for the quantification of the activity of undetected transcription factors, which is common in high-throughput single-cell transcriptomic data but to our knowledge not possible with SCENIC.

*SCORPION* enables the use of the same statistical techniques that account for population heterogeneity and are widely used in other areas of genomics data analysis by constructing very precise and highly comparable gene regulatory networks for each sample. We anticipate that *SCORPION* will be used not only to characterize molecular mechanisms driving phenotypes, but also to investigate a wide range of important questions in precision medicine, health, and biomedical research now that gene regulatory network perturbations have been shown to be effective at reproducing experimental results (70–72).

## Methods

### Generation of prior networks

To generate the unrefined regulatory networks that serve as prior for the message passing algorithm, we downloaded the promoter region coordinates for each gene from ENSEMBL. We then used TABIX (73) to retrieve the motif footprints and associated MOODS match scores located within 1000 bp prior to the transcription start site of each gene from Vierstra et al. (28). When multiple matches of the same transcription factor footprints were found, the highest value was retained for the study. The data on transcription factor protein-protein interactions and their associated scores were obtained from the STRING database version 11.5 (27).

### Benchmarking using synthetic data

BEELINE was used to conduct a systematic evaluation of cutting-edge algorithms for inferring single-cell gene regulatory networks (24). We used *SCORPION* and twelve other single-cell gene regulatory network inference algorithms on the GSD dataset, which is the largest dataset included in BEELINE and was generated from a curated Boolean model (74). These techniques include: GENIE3 (29), GRISLI (30), GRNBOOST2 (31), GRNVBEM (32), LEAP (33), PIDC (34), PPCOR (35), SCINGE (36), SCNS (37), SCODE (38), SCTENIFOLDNET (9), and SINCERITIES (39), SCRIBE (75) was excluded from the comparison due to compatibility issues. We processed the dataset using BEELINE’s uniform pipeline, which includes four steps: (1) data pre-processing, (2) docker container generation for *SCORPION* and the other 12 algorithms mentioned above, (3) parameter estimation, and (4) post-processing and evaluation. No information on TF-target relationships was provided to any of the algorithms we benchmarked *SCORPION* against throughout the analysis. We compared algorithms based on their average performance across seven different metrics: AUROC, AUPRC, computing time, level bias due to expression level, FBL, FFL, and MI motif structures identification. AUROC portrays a tested algorithm’s performance by presenting the trade-off between true positive rate 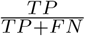 and false positive rate 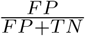 across different decision thresholds. AUPRC represents the area under the precision 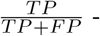 recall 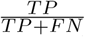 curve computed for different decision thresholds between 1 and 0 using, where *P*_*i*_ and *R*_*i*_ are the precision and recall at the *i*^th^ threshold. *TP* denotes true positive, *TN* denotes true negative, *FP* denotes false positive, and *FN* denotes false negative. The absolute value of the correlation between the average gene expression for each gene and its corresponding degree in the network was used to calculate the level bias due to expression level. FBL denotes feedback loops, while FFL denotes the feed-forward loop, a three-gene pattern composed of two input transcription factors, one of which regulates the other, both of which jointly regulate a target gene. Finally, MI stands for mutual interactions.

### Benchmarking using curated single-cell RNA-seq data

Count matrices for both experiments and conditions were downloaded from the GEO database with accession numbers GSM3477499, GSM347750, GSM5694433 and GSM5694434. Data was loaded into R using the build-in functions included in Seurat for this purpose (21). Two networks (one for the wild-type sample and one for the double knockout) were built for the Hnf4*αγ* experiment using *SCORPION* (under default parameters). The study was restricted to genes expressed in at least 5% of the cells in each sample. For the *DUX4* experiment, datasets were subject to quality control and integrated using Harmony (76, 77). Low dimensional representations and clustering of the data were generated using the top five dimensions returned by Harmony. 8 cell-like cells were annotated based on the expression of *ZSCAN4, DUXA, CCNA1* and *KDM4E* genes using the Nebulosa package (78). All cells from the wild-type sample were used to build a gene regulatory network that represented this group. Cells exhibiting the 8 cell-like markers in the *DUX4* overexpression group were used to generate a gene regulatory network representing them. The study was restricted to genes expressed in at least 5% of the cells in both samples. The information in the rows of the network representing the transcription factor of interest for each sample was contrasted to compare transcription factor activities among samples. The residuals of the linear model trained over the data in each case were used to assess the differences in the activity of the transcription factor over each gene. The residuals of the linear model and the marker genes provided by the PanglaoDB database were used to perform gene set enrichment analysis (79). Additional markers of the 8 cell-like cells were defined by differential expression using MAST after comparing the cluster expressing the known marker genes against all other cells (11).

### Colorectal cancer single-cell RNA-seq atlas construction

We collected multiple publicly available single-cell RNA-seq count matrices for human healthy adjacent tissue and different regions of colorectal tumors (see Data Availability). Datasets were loaded into R and combined into a single ‘Seurat’ object (21). Following that, data were subjected to quality control, with only cells with a library size of at least 1, 000 counts and falling within the 95 percent confidence interval of the prediction of the mitochondrial content ratio and detected genes in proportion to the cell’s library size being kept. We also removed all cells with mitochondrial proportions greater than 10% (76). We then used Seurat’s default functions and parameters to normalize, scale, and reduce the dimensionality of the data using Principal Component Analysis (PCA). Harmony was used for data integration (77). The top 50 dimensions returned by Harmony were used to generate the UMAP projections of the data. Cell clustering was carried out using Seurat’s built-in functions, default resolution, and Harmony embedding as the source for the nearest neighbor network construction. Clusters were annotated using Nebulosa (78) and the canonical markers provided by Qi et al. (80).

### Colorectal cancer gene regulatory network atlas construction

Using *SCORPION* under default parameters, we built a gene regulatory network for each cell type within each sample having at least 30 cells in the constructed colorectal cancer single-cell RNA-seq atlas. We only included genes that were expressed in more than five cells in each subsample. For each network, the sum of the activity of all transcription factors over each gene (in-degrees) was computed and assembled in a matrix. We used Principal Component Analysis to reduce the dimensionality of the data to the top 50 principal components. We used this data as input for the generation of the t-distributed Stochastic Neighbor Embedding (t-SNE) projection (81). Networks are color coded as their respective cell-type in the single-cell RNA-seq atlas.

### Modeling the regulatory differences that drive colorectal cancer progression

We selected the gene regulatory networks representing the epithelial cells (*EPCAM*^+^) of the different tumor regions (border, core and metastatic) and the healthy adjacent tissue. We modeled each edge weight representing the transcription factor - target gene interaction across the four different stages. We computed a *β* coefficient representing the average rate of change across each stage for each edge. The significance of the *β* coefficient was assigned using the F-distribution. Adjustment of the P-values for multiple testing was performed using False Discovery Rate (45).

### Comparison between right- and left-sided tumors gene regulatory networks

We selected the generated gene regulatory networks representing the epithelial cells from right- and left-sided tumors. For each network, we computed the (outdegrees) sum of all the activities for each transcription factor over all the genes. We then compared the outdegrees using the ttests function included in the Rfast package (82). P-values were adjusted for multiple testing using False Discovery Rate (45).

### Establishment of patient-derived tumor xenografts

The PDX models were derived in the same manner as described previously (83). The University of Texas at Austin and The University of Colorado Institutional Animal Care and Use Committee approved all animal procedures. Briefly, two to three *mm* pieces of colorectal tumor sample collected under IRB-approved protocol at the University of Texas Dell Medical School and the University of Colorado Cancer Center were engrafted onto the right and left hind flanks of 5 to 6-week-old Nu/Nu Mice (Envigo). Tumor volumes were measured by digital calipers every 3 to 4 days and were calculated by *V* = 0.52 × (*length* × *width*^2^). Mice were sacrificed when tumors reached 1.5 cm^3^ to further propagate the PDX model to the next generation or frozen as a viable tumor (RPMI media containing 10% FBS and 10% DMSO as a freezing media) in LN_2_ for long term storage. At the time of tumor harvest, a portion of the tumor was flash frozen in LN_2_ for RNA isolation and sequencing. RNA was isolated using PureLink kit (Thermo Fisher) following the manufacturer’s protocol. When the tumor specimen was abundant enough, a portion of the tissue sample was flash frozen, and RNA was isolated directly from that tissue. The RNA sample was outsourced to Novogene. US subsidiary and UC Davis Sequencing Center, Sacramento, CA for RNA QC, library preparation, and sequencing. Data obtained from Novogene as FASTQ files were subjected to further analysis.

### RNA-seq gene expression quantification

Gene expression from FASTQ files was quantified using STAR (84). The computed values for each PDX were loaded into R to generate the expression matrix. The t.test function was used to compare the expression levels of both (*ZNF350* and *NFKB2*) transcription factors.

## Data Availability

All of the data and code required to replicate the analysis as well as the figures and tables are available at https://github.com/dosorio/SCORPION. The *SCORPION* multi-platform stable package is available at https://CRAN.R-project.org/package=SCORPION. Versions under development are available at https://github.com/kuijjerlab/SCORPION. Patient-derived xenografts associated raw fastq files generated for this study are available upon request to Dr. S. Gail Eckhardt.

The following datasets were used to construct the colorectal cancer single-cell RNA-seq atlas used in this study: Lee et al. (85) accessible through GEO: GSE132465, and GSE144735. Qi et al. (80) accessible through GSA: [HRA000979](https://www.nature.com/articles/s41467-022-29366-6#Sec31), Qian et al. (86) accessible through ArrayExpress: E-MTAB-8107, and Che et al. (87) accessible through GEO: GSE178318.

## ACKNOWLEDGEMENTS

This work was supported by the Biomedical Research Computing Facility of the University of Texas at Austin. Figure 6A was created with BioRender.com.

## AUTHOR CONTRIBUTIONS

Conceptualization: DO, SGE and MLK. Methodology: DO, and MLK. Validation: DO, AC, UG, AS, TMP, CHL, WAM, SMB. Resources: DO, SSY, and MLK. Data Curation: DO. Writing - Original Draft: DO. Writing - Review & Editing: DO, SGE and MLK. Supervision: SSY, NS, and MLK. Project administration: SSY and MLK. Funding acquisition: AC, SGE, TMP, CHL, WAM, SSY, and MLK.

## Supplementary Figures

**Fig. S1.**
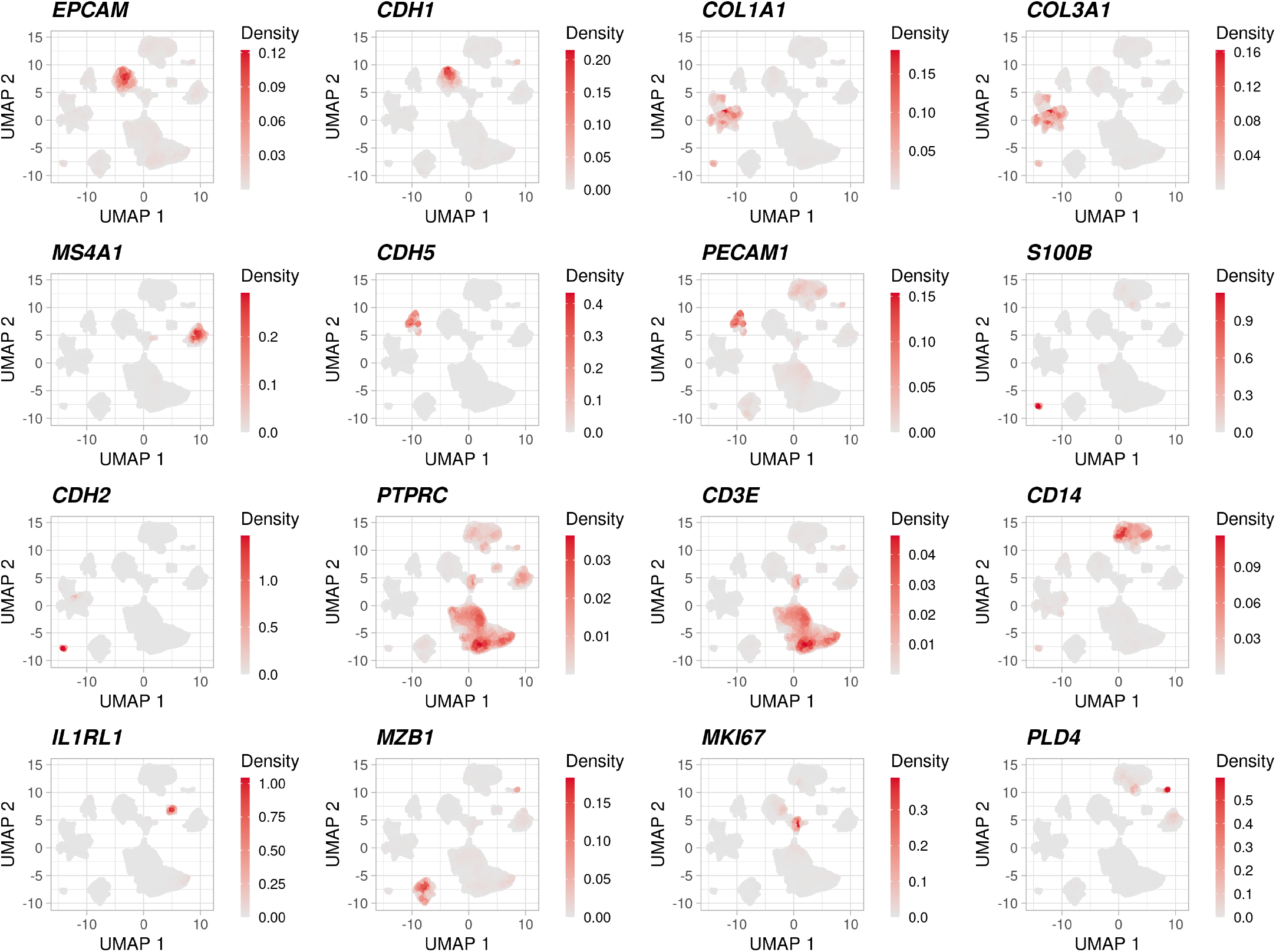
Distribution of canonical markers used to annotate cell types in the colorectal cancer single-cell RNA-seq atlas through the Nebulosa package

**Fig. S2.**
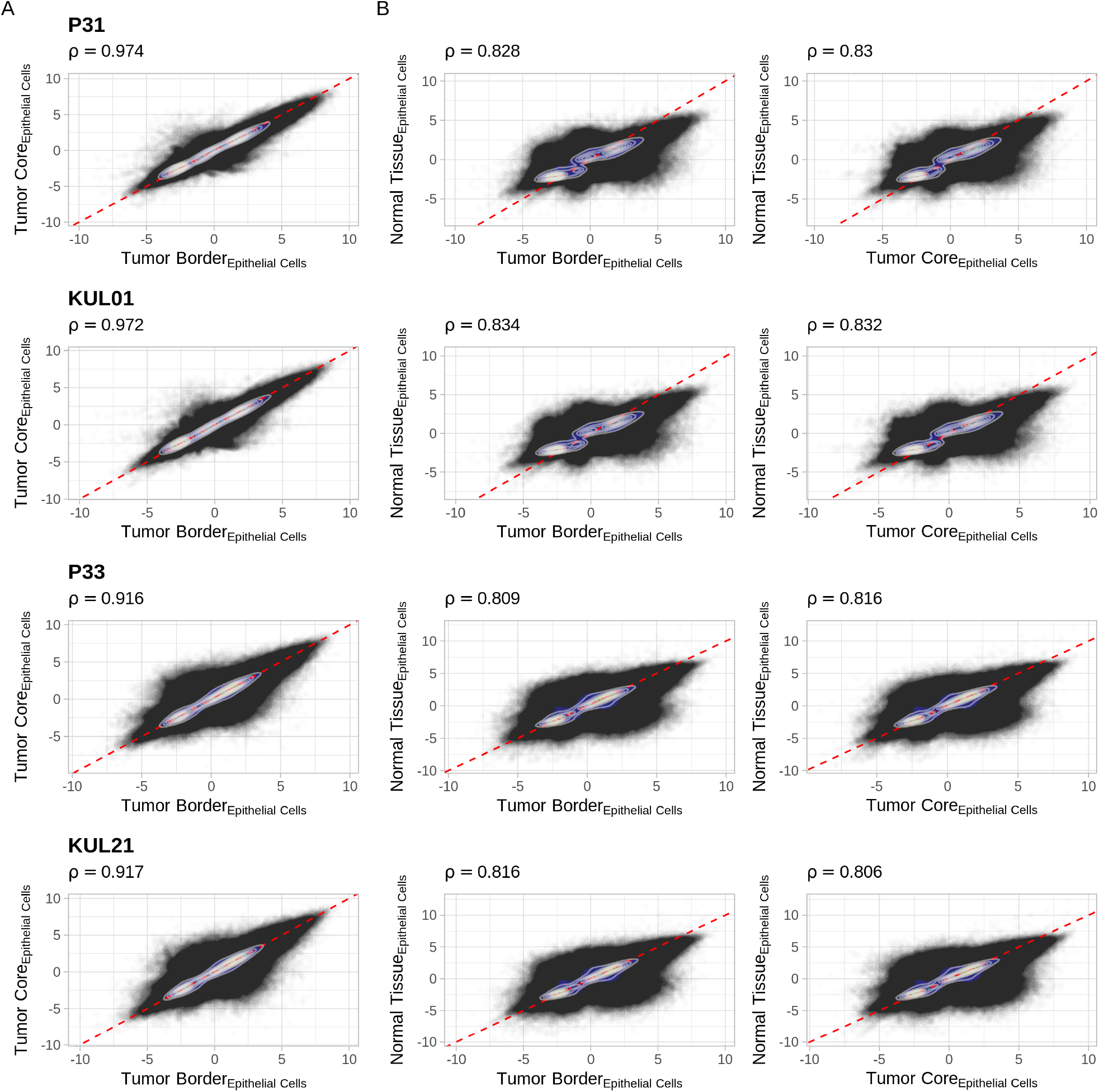
Similarity (Spearman correlation coefficient) between gene regulatory networks generated from the same donor for different tumor regions. **(A)** Similarity between gene regulatory networks generated for border and core of the tumor cancer tissue.**(B)** Similarity between gene regulatory networks generated for the core of the tumor, border of the tumor and healthy adjacent tissue.

## Supplementary Tables

**Table S1**. (CSV file) Numerical results from the BEELINE benchmark.

**Table S2**. (CSV file) Gene set enrichment analysis results for the *Hnf4α* knockout.

**Table S3**. (CSV file) Gene set enrichment analysis results for the *Hnf4γ* knockout.

**Table S4**. (CSV file) Identified marker genes and their associated fold-change and P-value for the 8-cells-like cells

**Table S5**. (CSV file) Gene set enrichment analysis results for the *DUX4* knockout.

**Table S6**. (CSV file) Metadata associated with the constructed single-cell RNA-seq atlas accounting for 200, 439 cells from different regions of colorectal cancer tumors and healthy adjacent tissue.

**Table S7**. (CSV file) Computed *β* coefficients and their associated P-value for each edge across the progression of colorectal cancer.

**Table S8**. (CSV file) Quantified gene expression levels from the generated PDX’s.

